# OligoY pipeline for full Y chromosome painting

**DOI:** 10.1101/2024.03.06.583648

**Authors:** Isabela Almeida, Henry Angel Bonilla Bruno, Mara Maria Lisboa Santana Pinheiro, Antonio Bernardo Carvalho, Maria Dulcetti Vibranovski

## Abstract

**Motivation:** The standard protocol for designing probes used in full chromosome fluorescent labeling experiments does not include repetitive sequences to avoid off-target hybridization. Due to the Y chromosome’s highly repetitive nature, most assembly nowadays still have heavily fragmented and incomplete Y sequences. Among these, the remaining non-repetitive sequences are insufficient to design probes and efficiently perform FISH Oligopaint assays, since they do not cover most regions of the chromosome. Ergo, cytogenetic studies with the Y are sparse, and analysis such as its function throughout the cell cycle and insights into its evolutionary history and relationships with other regions of the genome remain poorly studied.

**Results:** In this work, we introduce a new pipeline for designing FISH Oligopaint probes for the Y chromosome of any species of interest. OligoY pipeline uses open-source tools, enriches the amount of contigs assigned to the Y chromosome from the draft assembly, and effectively uses repetitive sequences unique to the target chromosome to design probes. Throughout all of its steps, the pipeline guarantees the user the autonomy to choose parameters, thus maximizing overall efficiency of cytogenetic experiments. After extensive in silico and *in situ* tests and validations with *Drosophila melanogaster*, we showed for the first time a pipeline for probe design that significantly increases previous Y chromosome staining with no off-target signal.

**Availability:** The pipeline is available at https://github.com/isabela42/OligoY.

## Introduction

Y chromosomes’ highly heterochromatic nature makes them prone to the accumulation of repetitive DNA (Skaletsky et al., 2003; Bachtrog, 2013), often resulting in their poor assembly in genome projects (Nishibuchi and Déjardin, 2017; Cechova, 2020; Tomaszkiewicz et al., 2017). A large proportion of the sequenced heterochromatin frequently remains as a collection of small, unmapped scaffolds, sometimes referred to as ”chromosome U” (Figure 1a) (Hoskins et al., 2002; Carvalho and Clark, 2013), which can be a result of repetitive reads collapse despite lacking unique positioning (Cechova, 2020). Some genome projects were specifically interested in improving the assembly of heterochromatic regions (Hoskins et al., 2002, 2007). Now, long-read technologies facilitated the detection of repeat-rich areas (Debladis et al., 2017), enabling telomere-to-telomere assemblies for all human chromosomes, Y inclusive, and also for a few other organisms (Wang et al., 2022; Nurk et al., 2022).

**Fig. 1:**
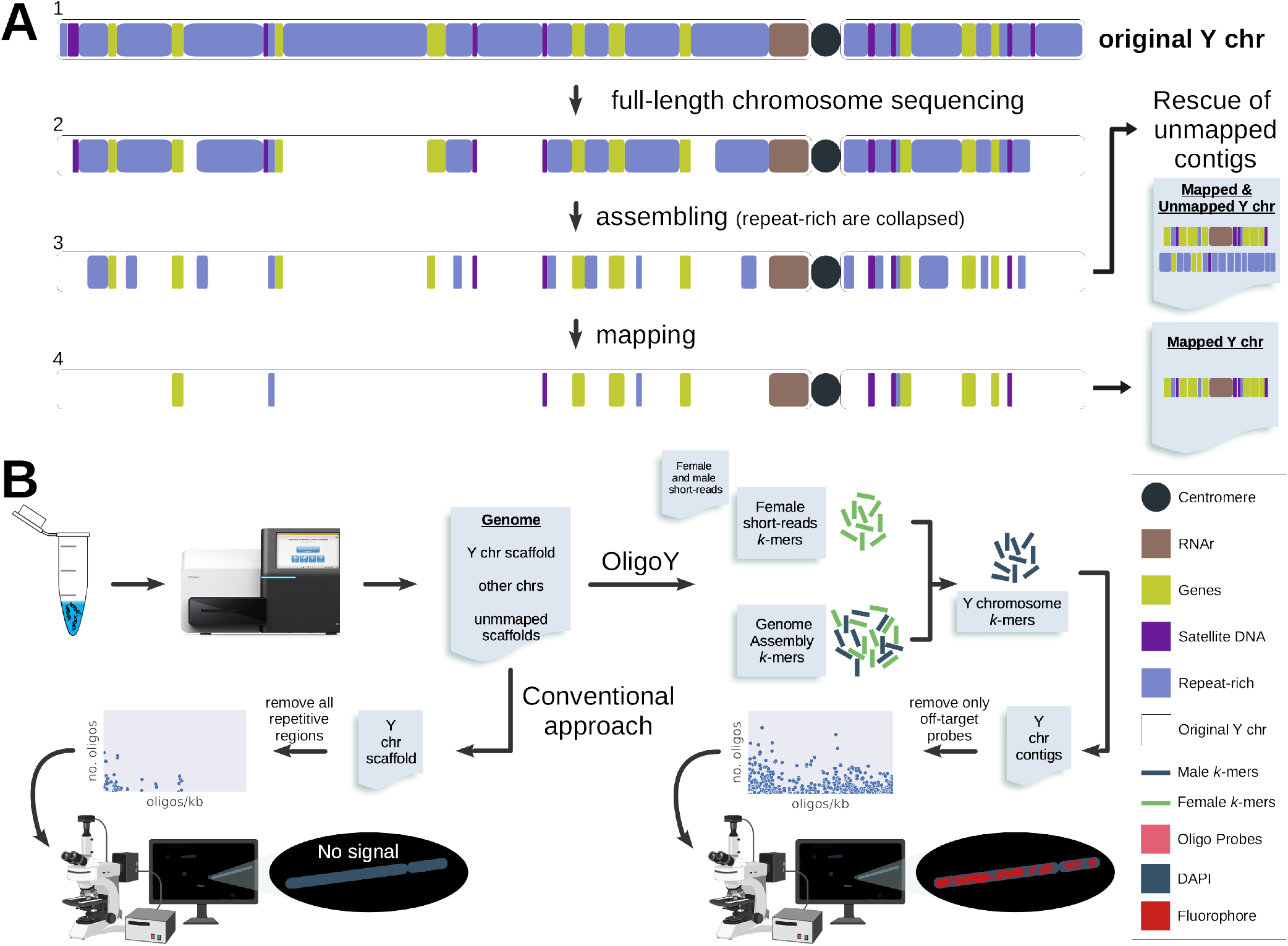
Structure, assembly, probe design and cytogenetic assays for Y chromosomes. A) The majority of the Y chromosome regions (1) are lost either during sequencing due to its haploid and repetitive nature (2), during assembly due to the collapse of repeat-rich reads (3) or during mapping, since only a few sequences, such as known genes, are mapped to the Y, with most remaining not assigned to a specific chromosome (unmapped); B) While conventional probe design for full chromosome fluorescent labeling assay does not use repetitive sequences to avoid off-target signals, resulting in fewer probes/kb and no fluorescence signals, OligoY pipeline includes all Y chromosome exclusive sequences, i.e. repetitive and non-repetitive, resulting in higher probe density and stronger signals in FISH Oligopaint assays.

Visualizing the Y chromosome completely would grant cytogeneticists the opportunity to examine how it interacts with other chromosomes and proteins during different cellular stages, as well as help to understand if and how the Y controls other chromosomes’ expression (Fingerhut et al., 2019; Chang and Larracuente, 2018). Among Y chromosome mapped regions, several conserved satellite-specific sequence motifs are found (Thakur et al., 2021). Because of this, individual satellite probes are typically used in cytological Y chromosome experiments (Jagannathan et al., 2017). For optimizing the signals detected during microscopy, such satellite probes must present motifs with a minimum of repetition, typically encompassing tandem satellite regions, while repeats dispersed across the target chromosome are not used (Bonaccorsi and Lohe, 1991). Since satellite probes only focus on a few areas and are shared with other chromosomes, they cannot be used to specifically study the Y chromosome in its entirety (Jagannathan et al., 2017).

The initial chromosome painting techniques in which individual chromosomes were isolated and used to make fluorescent DNA libraries were developed to overcome the limitations of satellite probes in covering entire chromosomes. To ensure the specificity of the hybridization process, a technique of suppression was employed (Pinkel et al., 1988; Lichter et al., 1988). Recently, many tools used whole genome sequences to create chromosome-specific single-copy probes, i.e. FISH Oligopaint probes, that combined hybridize along the entire target chromosome (e.g., Beliveau et al. (2018)). While this works elegantly for the autosomal and sexual X and Z chromosomes (Rosin et al., 2018; Nguyen et al., 2020; Rosin et al., 2021), for the Y/ W these approaches rely on the few remaining non-repetitive sparsely mapped sequences to avoid off-target hybridization. Ergo, using such tools to design probes for the Y chromosome in a conventional manner typically results in a quantity that is not worth doing cytological assays on (Figure 1b).

Here, employing open-source tools, we developed the OligoY pipeline for maximizing the Y chromosome covered area during efficient FISH Oligopaint assays. Using the YGS tool (Y chromosome Genome Scan, Carvalho and Clark (2013)), which compares male and female reads with an assembled genome, OligoY enriches the amount of contigs known to the Y chromosome by recovering the male exclusive sequences. Among these, there are numerous repetitive regions, and OligoY also uses those to design candidate FISH Oligopaint probes with the OligoMiner tool (Beliveau et al., 2018). In one of its filtering steps, OligoY presents a new approach for one of OligoMiner’s options, removing only those probes that effectively target other chromosomes, revoking the urgency to dismiss repeat-rich regions completely. Our test with *Drosophila melanogaster* results in higher expected probe density throughout the Y chromosome, along with signals during FISH Oligopaint assays combined with hybridization of DNA satellite probes that allow for proper Y chromosome observation.

## The pipeline

A genome assembly of the target species and its female and male reads are required for running the OligoY pipeline. There are two major steps to it: the Y chromosome contigs inference and the probe design (Figure 2a). In the first step, OligoY uses the YGS tool (Carvalho and Clark, 2013) to determine which sequences in the provided genome assembly are part of the Y chromosome. As a result, the Y chromosome has more known contigs and, more importantly, known nucleotide content, which can be used to design Oligopaint probes in the second major step using our novel OligoMiner tool approach (Beliveau et al., 2018). Users have the option to run the entire pipeline using three Bash scripts (Scripts A, B, and C; Supplementary Figure 1 and 2) or execute individual commands in the same order. Both methods yield identical results. Either approach yields identical results.

**Fig. 2:**
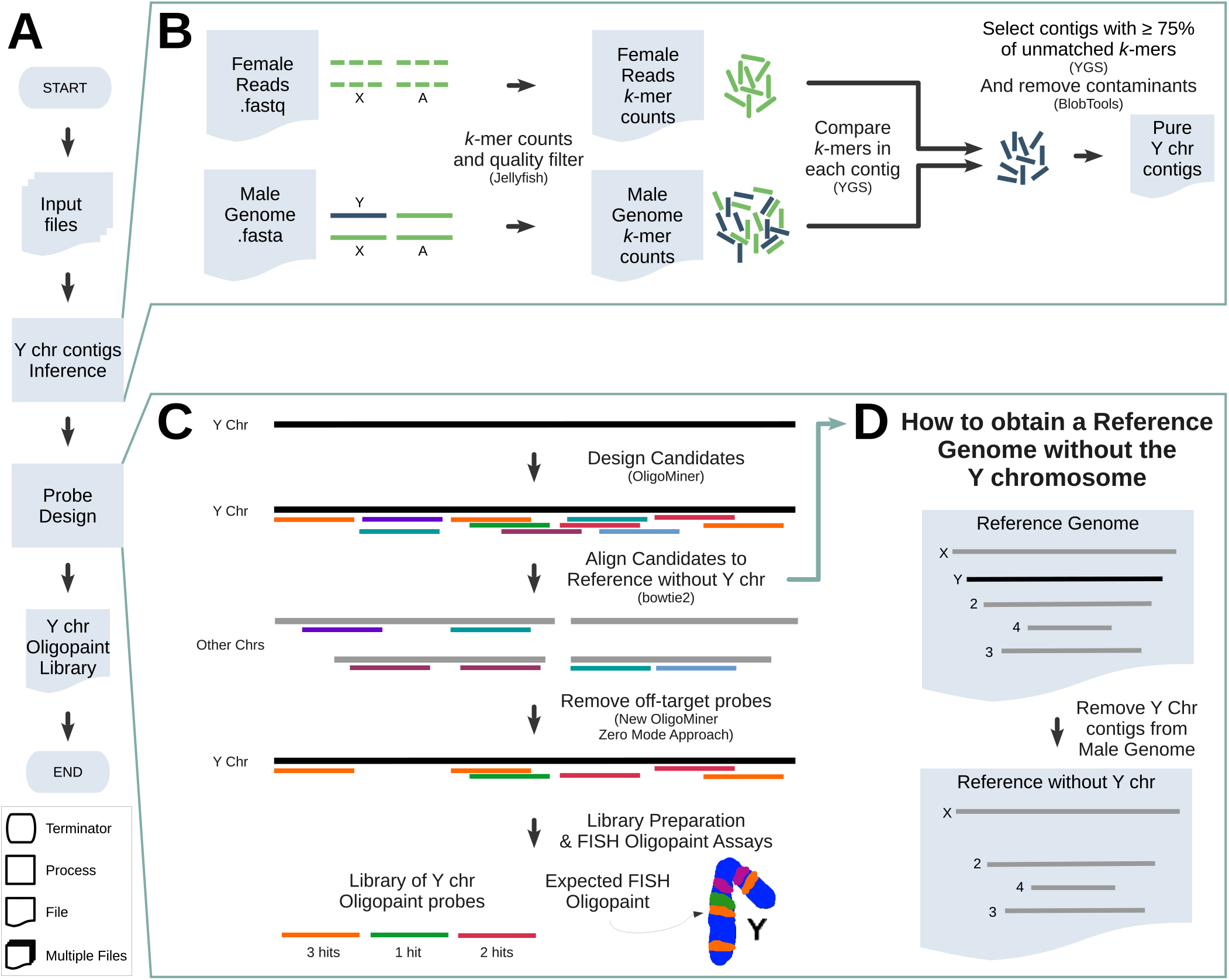
OligoY pipeline. A) OligoY in major steps in our pipeline; B) Y chromosome contigs inference step: YGS is implemented to compare *k*-mers from the male genome assembly and from the female sequencing reads. We recommend that selected Y chromosome contigs contain at least 75% of valid single-copy *k*-mers (VSCK) unmatched by female reads (PVSCUK) and meet a minimum threshold of 20 VSCK; C) Probe design step: candidate probes are designed and filtered against a reference genome without the Y chromosome sequences; with our novel OligoMiner’s Zero Mode (ZM) approach, OligoY pipeline only removes probes that effectively target other chromosomes, producing a library with both single and multiple hits, all exclusively Y probes; D) By removing the Y chromosome inferred contigs from the original genome file we have the reference genome without the Y chromosome.

In the first main step of OligoY, input files are independently preprocessed using Jellyfish (Marçais and Kingsford, 2011) to create high-quality trimmed hash tables (Figure 2b). Next, the pipeline runs YGS’s bit-vector converter, which calls Jellyfish internally to produce a dump bit-vector without rare *k*-mers. Importantly, the dump bit-vector created from the genome assembly must only contain repetitive *k*-mers (lower-count *>*1). Next, YGS is executed in its final run mode. At this point, YGS will first create another bit-vector for the genome with all of its *k*-mers and then all the bit-vectors previously generated are compared to each other to create internal ones for each possible combination. At the end of its analysis, YGS generates a report with a series of metrics for each genome sequence, including the proportion of valid single-copy *k*-mers (VSCK) and the proportion of *k*-mers that are unmatched by female reads (PVSCUK). These are an important metric used by OligoY to ascertain which sequences belong to the Y chromosome when they fall within a certain range (Figure 2b). Indispensably, YGS also removes genome contaminants by comparing the different bit-vectors to the one generated from male reads. These commands are comprised within Script A and its key output is the Y chromosome FASTA file. Following that, the OligoY pipeline includes a contaminant analysis step (Script B) using BlobTools (Laetsch and Blaxter, 2017; Vanderlinde et al., 2018), which avoids inferring any male-specific contaminated sequence to the Y chromosome and designing Oligopaint probes to it. For this, we call its key output the pure Y chromosome FASTA file.

In the second main step of OligoY (Script C), overlapping candidate probes are designed from the concatenated pure Y chromosome inferred sequences utilizing OligoMiner (Figure 2c). Employing our new OligoMiner approach, these candidate probes are aligned against the input genome assembly without the Y chromosome inferred sequences (Figure 2d) using bowtie2 (Langmead and Salzberg, 2012). Next, OligoMiner’s filter of predicted target number of alignments can be used with the Zero Mode (ZM) option, removing all the probes that effectively aligned outside the Y (Filtering level 1). This is achieved by keeping those probes that are aligned zero times in the provided genome – a genome assembly without the Y chromosome. Therefore, only exclusive Y chromosome probes are kept. This strategy allows OligoY to maintain Y chromosome probes whether they appear once on the chromosome or repeatedly. Due to the complexity of this chromosome, additional OligoMiner filtering steps are also used, such as the elimination of probes that have an abundance of *k*-mers in other chromosomes (Filtering level 2) and of those that are likely to form secondary structures (Filtering level 3). All potential probe regions can be identified by using overlapping probes, but synthesized probe overlapping is detrimental to *in situ* experiments, such as the hybridization procedure. As a result, the OligoY pipeline includes an in-house Python program for removing overlapping probes from those that have passed all previous filtering steps (noOverlap.py, Filtering Level 4). We also implemented an in-house Python program to retain only the first occurrence of a probe, avoiding redundancy, since multiple syntheses of the same probe are unnecessary and only matter for probe density estimations (nonRedundant.py, Filtering level 5). Finally, OligoY has a sixth filtering level that selects probes with no alignments to female reads (converted to genome sequence with seqtk by Li – available at https://github.com/lh3/seqtk), removing the possibility of probes in the final library aligning outside the Y chromosome.

## Pipeline development and validation

*Drosophila melanogaster* provided us with a highly repetitive Y chromosome and incomplete genome assemblies that are likely to mimic the situations faced by users of the OligoY pipeline. Namely, the ones here mentioned as Illumina (Gutzwiller et al., 2015), Sanger (Hoskins et al., 2002) and Release 6 (Hoskins et al., 2015) were the most fragmented draft genomes we used, as opposed to Chang and Larracuente (2018) and the Falcon genome (Kim et al., 2014) (Supplementary Table 1).

Our OligoY technique significantly outperformed the default OligoMiner analysis for the *Drosophila melanogaster* Y chromosome, generating a markedly higher number of probes. While OligoMiner produced 587 to 1,757 probes across varying stringency levels, corresponding to lengths and experimental conditions that range from more rigorous to more permissive settings (Beliveau et al., 2018), OligoY developed 6,428 to 19,007 probes under comparable conditions using YGS-derived sequences from the same input as (Beliveau et al., 2018), i.e. Release 6 (Figure 3a, Supplementary Tables 11 and 12). To dissect the precise processes responsible for probe yield enhancement, our in silico tests were designed to evaluate the outcomes of each major step within the OligoY approach: Y chromosome contig inference and probe design.

**Fig. 3:**
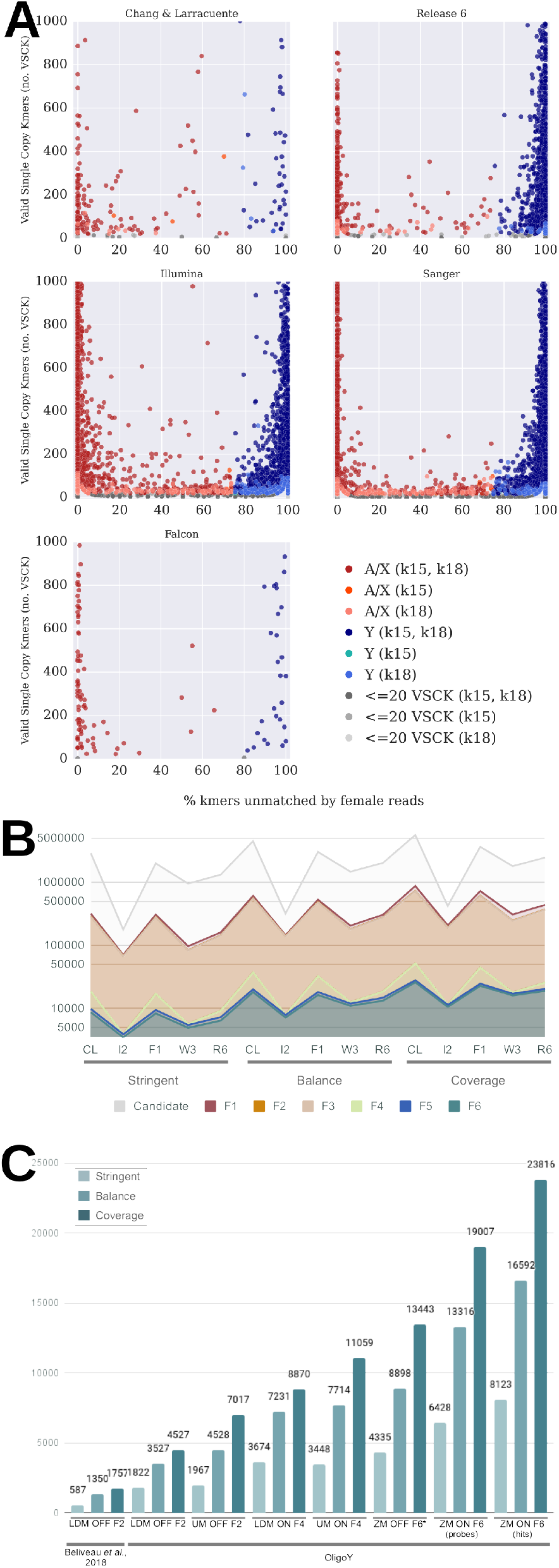
Parameter evaluation A) There is a substantial overlap in between the inferred sequences when running YGS with *k*-mer values 15 and 18, regardless of input draft genome; B) Loss of probes throughout OligoY filtering steps (log scale, F1 to F6 represent filtering levels 1 to 6); C) Number of probes designed by OligoY with YGS-derived Release 6 Y chromosome sequences compared to the ones designed by Beliveau et al. (2018) (first three bars to the left). ^*^For ZM, when overlapping probes were not allowed (OFF), the resulting F6 probes were not filtered for level 4, which removes overlapping proves.

### *In silico* tests improve nt content for inferred Y chromosome

Our tests on Y chromosome contig inference demonstrated a consistent pattern across the most and less fragmented input genomes. Regardless of *k*-mer values employed (15 or 18), we consistently observed more distinct contigs being inferred for each individual Y chromosome group of sequences for Illumina (I2), Sanger (W3), and Release 6 (R6) assemblies, which had the shortest nucleotide content sizes among the tested ones. In fact, Illumina’s estimated Y chromosomal sequences were found to be as low as less than 1.5 megabases (Mb). Conversely, when using Falcon (F1) and Chang and Larracuente (CL) input genomes, the less fragmented ones, we identified up to 13.8 Mb of Y chromosome sequences (Figure 3a, Table 1, Supplementary Tables 02 and 03).

**Table 1.** Technical results of YGS tests.

| Assembly | $k$-mer value 15 | | $k$-mer value 18 | | Common<sup>1</sup> | |
| --- | --- | --- | --- | --- | --- | --- |
|  | # | Mb | # | Mb | # | Mb |
| CL | 40 | 13.506146 | 44 | 13.789009 | 40 | 13.506146 |
| I2 | 1897 | 1.153488 | 2964 | 1.299917 | 1887 | 1.150502 |
| F1 | 66 | 8.80499 | 66 | 8.80499 | 66 | 8.80499 |
| W3 | 2659 | 5.507152 | 3449 | 6.185429 | 2654 | 5.50321 |
| R6 | 769 | 6.567302 | 821 | 6.650289 | 768 | 6.566277 |

Number (#) of contigs and total size in megabases (Mb) are given
<sup>1</sup>Sequences inferred by both $k$ -mer values

Over half of the sequences in Release 6 are unmapped contigs (Supplementary Table 04), accounting for 3.05 Mb of genetic material. The Y chromosome in this assembly was made up of a single scaffold and 199 contig sequences, amassing a total of 4.53 Mb (Hoskins et al., 2015). While using this assembly as in input, YGS implementation in our pipeline facilitated the inference of numerous unmapped contigs as part of the Y chromosome, thereby augmenting its known sequences to a minimum of 769 contigs (6.57 Mb, using a *k*-mer of 15) and a maximum of 821 contigs (6.65 Mb, using a *k*-mer of 18; Table 1, Supplementary Table 03).

Importantly, 1.15 Mb of sequences (16 contigs) assembled by Chang and Larracuente (2018) from the Y chromosome were not inferred as such using YGS with either *k*-mer value 15 or 18. One of these was identified as a contamination, and the remaining did not meet our minimum threshold for the number VSCK (20). Additionally, our YGS analysis selected two additional sequences, each with at least 97.3% of valid *k*-mers that were not present among the female reads. While Chang and Larracuente (2018) relied on an extensively detailed approach with a series of manual curations to enrich the known sequences for the Y chromosome of *Drosophila melanogaster*, OligoY users can achieve similar results with significantly less effort and meticulousness because we have incorporated YGS into it.

Notable for all the conducted tests, *k*-mer value 18 didn’t show significant advantages over *k*-mer value 15. Hence, we advise using *k*-mer 15 for smaller genomes (e.g., invertebrates) and 18 for larger ones (e.g., vertebrates), aligning with Carvalho and Clark (2013) recommendation. In addition, we confirmed no contaminated sequences in the inferred Y chromosome data (Supplementary Tables 05 and 06). Only sequences classified under Arthropoda or Eukaryota taxa were retained, ensuring no wrongful probe design.

### Designing probes with a density of over 1 probe/kb

The success of OligoY in achieving a much denser coverage of the target chromosome is intricately connected to four key features: (i) to design probes from a YGS-derived inputs, i.e. more extensive nucleotide sequence compared to the initial inputs; (ii) extensive filtering steps to guarantee no contaminants and off-target probes are kept; (iii) designing candidate probes that meet the desired criteria with the overlap option, increasing the chances of retaining signals from regions that might otherwise be discarded; and (iv) allowing multiple hits in the target chromosome to be kept.

Previously, Beliveau et al. (2018) designed Y chromosome probes for the Release 6 genome targeting different settings from strict to permissive, resulting in oligopools of 587 (stringent), 1350 (balance) and 1,757 (coverage) probes. They used OligoMiner’s linear discriminant analysis (LDA) mode and kmerFilter.py to remove probes with abundant *k*-mers. When solely changing the input target chromosomes to a YGS-derived one, we were able to design 1,822, 3,527 and 4,527 for each of those respective parameter conditions (Supplementary Tables 11 and 12). This shows that by implementing YGS in our pipeline and inputting a more extensive nucleotide sequence, we more than double the number of target probes that will compose the oligopaint library.

Regardless of the input genome and experimental conditions we tested, OligoY produces numerous candidate probes that undergo extensive filtering. Most of them are lost in the first three filtering steps, highlighting their effectiveness in ensuring that only non-off-target probes are kept in the final pool, even though probes with multiple hits inside the Y chromosome are still retained (Figure 3b, Supplementary Table 11). Moreover, while the final filtering steps remove just a few probes (Figure 3b) they assure that (i) final probes are not overlapping sequences, which would negatively affect FISH assays, (ii) only one copy of each probe is kept, since candidates with multiple hits are designed from multiple matching target regions, and (iii) none of these probes align against the female reads, which could have been abundant fragments not assembled for some reason.

To enhance the likelihood of preserving signals from regions for which candidate probes were filtered out, we used the native overlapping option from OligoMiner. Comparing to Beliveau et al. (2018) use of the LDA mode, we now designed from 3,674 (stringent) to 8,870 (coverage) probes with the YGS-derived Release 6 Y chromosome sequences (Supplementary Tables 11 and 12). Despite the LDA mode keeping an initial larger number of probes for simulating the number of hybridization in the reference genome, when applying further filtering steps most of the probes with more than one hit are removed. Using the same input and OligoMiner’s unique mode (UM), our tests resulted in 3,122, 7,101 and 10,415 probes (stringent, balance and coverage parameters), over 5x the number of probes designed by Beliveau et al. (2018).

With the feature in our pipeline that allows probes with multiple hits to be kept, we designed up to 26,323 probes with 48,497 hits on the Y chromosome (coverage and *k*-mer 18 parameters with YGS-derived Chang and Larracuente’s genome). The same parameters and input with the standard OligoMiner mode yielded only 11,593 (UM) and 9,423 probes (LDA, Figure 3c). For the YGS-derived Release 6 Y chromosome sequences, the same settings yielded 19,264 probes with 24,075 hits as compared to the UM 10,436 and LDA mode 8,352 single-hit probes.

We validated YGS-derived Chang and Larracuente and Release 6 oligopaint probe libraries’ produced from the stringent and balanced settings from Y chromosome sequences commonly inferred with both tested *k*-mer values. Both inputs achieved a probe density of 1.2 to 3.6 probes/kb across these settings (Supplementary Tables 11 and 13). Yet, Chang and Larracuente’s sets of probes has roughly double the hits and were 30% larger than those of Release 6, despite having twice as many sequences.

### *Drosophila melanogaster in situ* tests

The performance of YGS-derived Chang and Larracuente’s (CL) and Release 6 (R6) oligopaint probe libraries’ produced from the stringent and balanced settings was assessed using FISH Oligopaint assays (CL-stringent, CL-balance, R6-stringent, R6-balance). Despite the highly repetitive nature of the Y chromosome, all four oligopaint libraries demonstrated remarkable specificity, with no evidence of off-target hybridization in other chromosomes (Figure 4c-f, Supplementary Figures 4-7). Interestingly, adopting more stringent length and experimental conditions to design Y oligopaint probes with OligoY is not a requirement. In fact, the more relaxed parameters we examined maintained target specificity (Figures 4c and 4e, Supplementary Figures 4 and 6). Our microscopy image analysis suggests a correlation between the number of oligos per library and fluorescence intensity (Supplementary Tables 11 and 13). In this context, CL libraries fluoresce more than R6 libraries, while the balance libraries fluoresce more than the stringent ones. Moreover, the CL-balance library displayed two additional labels near the ends of each arm – between bands *∼*2-4 and bands *∼*24-25 on the long and short arms, respectively (Figure 4a,b and Supplementary Figure 4). Bands 2–4 labels were also present in the R6-balance library at a lower fluorescence intensity (Supplementary Figure 6), and these findings support our suggestion that more permissive sets of length and experimental conditions can be used to design probes for the Y chromosome. Particularly to *Drosophila melanogaster*, our findings indicate that the CL-balance library represents the optimal choice for Y chromosome labeling, offering a compelling balance between specificity and broader coverage.

**Fig. 4:**
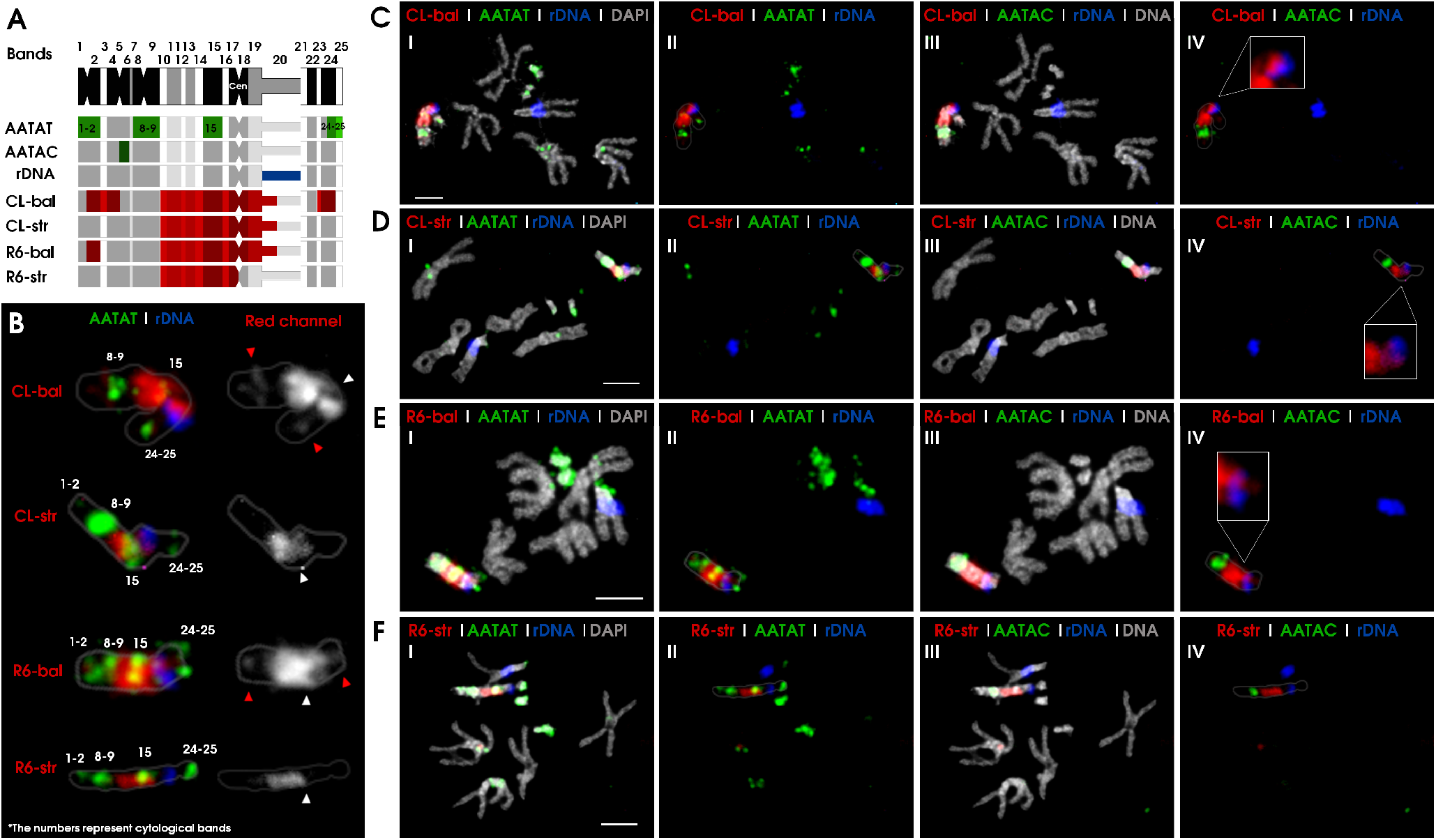
FISH oligopaint on larval neuroblast chromosome spreads from *Drosophila melanogaster* (ISO1 strain) for CL and R6 assemblies. A) Schematic representation of the Y chromosome showing 25 cytological bands and the approximate cytological location of the satellite probes (AATAT, AATAC) (Bonaccorsi and Lohe, 1991), ribosomal DNA (rDNA) (Chang and Larracuente, 2018) and the oligopaint libraries (CL-balance, CL-stringent, R6-balance, R6-stringent). Cen: centromere. B) Comparison of the labeling patterns observed in all four oligopaint libraries on the Y chromosome. The AATAT and rDNA probe positions are reference points for contextualizing the oligopaint labeling. Oligopaint labels in the red channel. Red arrowheads show the extra labels of balance libraries (CL-bal, R6-bal). The centromeres are indicated by white arrowheads. Images with FISH experiments for: C) CL-balance library, D) CL-stringent library, E) R6-balance library and F) R6-stringent library. I and III show the labeling of three probes (oligopaint, satellite and rDNA) combined with DAPI, while II and IV show only the three probes (colors specified on panel text). The zoom-in boxes in IV reveal the violet color formed by partial overlapping between oligopaints (red) and rDNA (blue). Note that oligopaints libraries (red) exclusively label the Y chromosome. CL-bal (Cy5: red), CL-str (Cy5: red), R6-bal (Cy5: red), R6-str (ATTO488: pseudocolor in red), AATAT (avidin-FITC: green), AATAC (avidin-FITC: green), rDNA (antiDIG-rhodamine: pseudocolor in blue) and DAPI (pseudocolor in gray). Bar, 2.5 mm.

Collectively, three libraries (CL-balance, CL-stringent, R6-balance) consistently tagged two areas on both sides of the Y chromosomal centromere (Figure 4a-e) while the R6-stringent library exclusively hits a portion of the long arm (Figure 4b,f and Supplementary Figure 7). Using two known satellite probes positions as reference (AATAC and AATAT, Bonaccorsi and Lohe (1991)) we demonstrated that our OligoY probes target Y chromosome cytological bands 10 to 20, including the pericentromere and the ribosomal DNA (rDNA) region (Chang and Larracuente, 2018) (Figure 4a). Our oligopaint libraries outperformed the tested satellite probes in both coverage and specificity: while the Y chromosome has been extensively and exclusively stained with our OligoY probes, AATAT satellite probe targets four much smaller chromosome sites, predominantly mapping to chromosome 4 and to a lesser extent targeting chromosomes 3 and X (Figure 4c-f, I and II). Moreover, although the AATAC satellite probe specifically labels the Y chromosome, this is limited to the small region of band 6 on the long arm (Figure 4c-f, III and IV).

Despite the presence of rDNA in *Drosophila melanogaster* on both sex chromosomes, X and Y (Jagannathan et al., 2017), our OligoY Oligopaint libraries selectively target Y-linked rDNA, showcasing our method’s ability to retrieve Y-linked probes despite their original sequences repetitiveness. The target rDNA region was effectively observed from the overlap between the signals from a rDNA probe (blue channel) and the rDNA region from CL libraries (red channel), resulting in the emergence of a violet staining in the Y chromosome band 20 area (Figure 4c-IV,d-IV).

## Conclusion

Designed for comprehensive Y chromosome cytology studies, OligoY pipeline finds its ideal testing ground in the Y chromosome of *Drosophila melanogaster*. We recognized the importance of incorporating YGS into our pipeline to enhance the available nucleotide content for probe design, considering that this tool offers a robust methodology and comprehensive insights into each of the genome sequences. Combined with it, OligoMiner suite allows its users to set a broad range of parameters and comes with many filtering steps and designing options. Here, we tested various combinations of these, finally pursuing a new approach with OligoMiner’s ZM filter, which in OligoY allows for probes with multiple hits in the target chromosome to be kept. This increases not only the final number of probes, but also the expected probe density throughout the Y chromosome.

The application of the OligoY pipeline is evident in our FISH Oligopaint assays, where precise targeting of the Y chromosome was achieved without any off-target hits. OligoY’s *in situ* validation experiments led to extensive coverage of the Y chromosome arms, including previously sparsely characterized heterochromatic regions in *Drosophila melanogaster*. Previously, we were limited to observing specific heterochromatic regions, e.g., h6 (AATAC) and h1-2, 8-9, 15, 24 (AATAT), leaving substantial gaps in our comprehension of Y chromosome structure. OligoY’s implementation resulted in the staining of numerous additional regions, significantly enhancing our capacity to comprehensively explore the Y chromosome’s architecture. This includes regions that were previously elusive, such as the *∼*2-4 and *∼*24-25 bands.

Our validations, restricted to larval metaphases due to the absence of Y polytene chromosomes in our model, hint at the potential for expanded use of OligoY in experiments across different and less condensed cell cycle phases. Furthermore, our Y chromosome labeling was achieved using oligos designed based on a set of known sequences that were 6.57 Mb (R6) and 13.5 Mb (CL) in size out of an estimated size that exceeds 40 Mb. While it is possible that some repetitive sequences within our oligos may correspond to previously unsequenced or unassembled regions, our results remain significant in their coverage of this complex chromosome.

## Materials and Methods

### Input files

We used five *Drosophila melanogaster* draft assemblies created with different sequencing technologies and methodologies (Hoskins et al., 2002; Kim et al., 2014; Hoskins et al., 2015; Gutzwiller et al., 2015; Carvalho et al., 2016; Chang and Larracuente, 2018)(Supplementary Table 1) in addition to Pacific Biosciences male (Kim et al., 2014) and Illumina female (Goldstein, 2016) short-reads from the ISO1 strain.

### Bioinformatics tools and packages

We used the following open-source tools: YGS (Carvalho and Clark, 2013); Jellyfish (Marçais and Kingsford, 2011); BlobTools (Laetsch and Blaxter, 2017); BLASTn 2.2.31+ and BLASTx (Altschul et al., 1990; Buchfink et al., 2014); OligoMiner (Beliveau et al., 2018); and bowtie2 (Langmead and Salzberg, 2012). OligoMiner requires: NUPACK package (Dirks and Pierce, 2003, 2004; Dirks et al., 2007) and Jellyfish to be installed on the same PATH, as well as scikit-learn 0.17+ package (Pedregosa et al., 2011). OligoY uses seqtk (available at https://github.com/lh3/seqtk). We converted OligoMiner’s structureCheck.py to Python 3+ using 2to3 API (available at https://docs.python.org/3/library/2to3.html), due to library availability.

### OS and Languages

High-performance computer specifications: Ubuntu 16.04.7 LTS (GNU/Linux 4.15.0-041500-generic x86 64); 17 T HD; 300 GB RAM; 40 CPUs; 2133 MHz max memory; 3100 MHz max CPU; IIntel^®^ Xeon^®^ CPU E5-2630 v4 @ 2.20 GHz. Our pipeline integrates the currently available open-source tools in Bash scripts using basic UNIX commands, and we developed Python 2.7+ programs for new filtering steps (noOverlap.py nonRedundant.py).

### Probe design through pipeline development

*k*-mer values of 15 and 18 were used for sequence inference on each genome input with other fixed parameters (Supplementary Table 2). After contamination removal (Supplementary Tables 05 and 06), these groups of inferred Y chromosome sequences were used to determine the best OligoMiner probe design mode (UM, LDA mode, and ZM) for OligoY. Three parameter sets (Supplementary Table 08) with and without overlap were tested for probe design.

### Library preparation

Our library was prepared to be amplified with the In Vitro Transcription – Reverse Transcription protocol (Beliveau et al., 2017). Oligonucleotides contain the following regions in the given order: forward priming, reverse transcription priming, target oligopaint probe and reverse priming regions, all sequences with 5’-3’ polarity (Supplementary Figure 3a). Priming sequences were selected based on previous work (Beliveau et al., 2015; Chen et al., 2015) and recommendations (Boettiger et al., 2016; Chen et al., 2015; Moffitt and Zhuang, 2016). We avoided guanine quadruple with the T7 promoter’s GGG-terminus on the synthesized reverse primer by ensuring no priming region had guanine on its 5’. We aligned 1) prospective priming regions against each other, removing those with a similarity ≥10 base pairs (bP); 2) prospective priming regions against the T7 promoter 3’ terminal, removing those with a similarity ≥5 bP; 3) prospective forward and reverse priming regions against all designed probes, removing those with a similarity ≥12 bP; 4) prospective reverse transcription priming regions against all the designed probes, removing those with a similarity ≥20 bP; 5) the entire oligonucleotide sequence against the individual priming regions after adding specific priming regions (Supplementary Table 09) to corresponding probes, ensuring no new similarity points were created (Altschul et al., 1990). In our library, all oligonucleotides had the same reverse priming region; the forward priming region changed according to the assembly from which the probes were designed; the reverse transcription priming region changed according to the set of parameters for probe design. To accommodate oligopaint probes in a single 90K microarray chip by CustomArray Inc. (a member of GenScript), only stringent and balance probes were synthesized. Additionally, any exact copies of oligopaint probes that were designed from two different assembly genomes were synthesized only once and contained both forward priming regions, allowing for individual group amplification (Supplementary Table 09 and Supplementary Figure 3b). GenScript ensured uniform size by adding random nucleotide bases at the 3’ end of the oligonucleotides.

### *In situ* validation

Real-Time PCR followed by T7 amplification and reverse transcription (Beliveau et al., 2017) was used to select and produce each individual group of probes within the synthesized oligonucleotide library (Supplementary Figure 3a,b). The used forward primer has the same sequence and polarity as the forward priming region on the oligonucleotide probe; the reverse primer starts with the T7 promoter followed by the reverse complement of the reverse priming region on the oligonucleotide probe (5-3’ polarity); the reverse transcription primer has the same sequence and polarity as the reverse transcription priming region on the oligonucleotide probe. Primers were synthesized by Sigma Aldrich/Merck (Supplementary Table 10). We used *Drosophila melanogaster* strain ISO1 (Clark et al., 2006), maintained in a hot chamber at 22 ^*°*^C and 12h:12h light/dark. We used brains of male third instar larvae to produce slides with metaphasic mitotic chromosomes by selecting male specimens with large translucent gonads on the dorsal part of their bodies (Maimon and Gilboa, 2011; Park et al., 2018) (Supplementary Figure 3c). Mitotic chromosome spreads were obtained with Mahadevaraju et al. (2021) protocol with some modifications: 10 to 12 brains extracted, washed twice with 100 *µ*l of 1X PBS, incubated in 100 *µ*l of KCL (0.075 mM) for 5min. Next, 30 *µ*l of KCl was removed and 30 *µ*l of fresh Carnoy Fixation Solution (3 Absolute Ethanol : 1 Acetic Acid) was added to the solution, and incubated for 5min. Then, the liquid solution was completely removed, and the brains were immersed in 100 *µ*l of Carnoy Fixation Solution for another 5min. Then, the brains were transferred to a microtube with 75 *µ*l of Acetic Acid Fixing Solution (60%). Finally, 20 *µ*l of the homogenized brain solution was spread on slides preheated to 40 ^*°*^C. Slides were kept at -20 ^*°*^C. FISH Oligopaint assays were performed according to Nguyen and Joyce (2019) protocol. CL-Balance, CL-stringent, R6-balance, R6-stringent oligopaints, (AATAT)6-biotin, (AATAC)6-biotin (Sigma Aldrich/Merck) and rDNA-digoxigenin probes were incubated at 37 ^*°*^C overnight in a humid chamber. (AATAT)6 and rDNA probes were incubated at 19 ^*°*^C. Images were acquired in a Zeiss Axioplan 2 microscope equipped with AxioCam MRm and a 100X oil immersion objective lens (NA = 1.4) (Carl Zeiss, Inc., Jena, Germany) and Isis V5.2 (MetaSystems GmbH) software. We processd images with GIMP 2.10.20 (The GIMP Development Team, 2012) and ImageJ (Schneider et al., 2012). Image resolution for the DAPI channel was improved using deconvolution. Then, point-spread function (PFS) was calculated using the Born and Wolf 3D optical model implemented in the plugging PFS generator Kirshner et al. (2012) for ImageJ. The model was run with settings according to our microscope (1.4 for numerical aperture NA, 67.5 for pixel size XY), 160 nm for Z-step (25 step) and 461 nm of emission wavelength (DAPI). Next, PFS was used to run deconvolutionLab2 with the Richardson-Lucy algorithm with 50 iterations in ImageJ (Sage et al., 2017) (Supplemental Figure 8).

## Author contributions statement

I.A., A.B.C. and M.D.V. conceived the project. I.A. designed and directed the study, performed all dry-lab analyses, developed the pipeline, critically reviewed the computational code, created and maintains the GitHub repository. H.A.B.B. implemented the FISH Oligopaint protocol in the laboratory and performed the majority of wet-lab experiments. M.M.L.S.P. produced male slides. A.B.C. provided genome assemblies. I.A. and M.D.V interpreted the data and co-wrote the manuscript with input from all authors.

## Acknowledgments

This work was supported by: Fundação de Amparo à Pesquisa do Estado de São Paulo (M.D.V.: 2015/20844-4, I.A.: 2019/14878-4, H.A.B.B.: 2019/10559-1); Wellcome Trust (A.B.C.: 207486/Z/17/Z); and Fundação Carlos Chagas Filho de Amparo à Pesquisa do Estado do Rio de Janeiro (A.B.C.: CNE2018).

